# Chemogenetic activation of midline thalamic nuclei fails to ameliorate memory deficits in two mouse models of Alzheimer’s disease

**DOI:** 10.1101/2021.06.30.450500

**Authors:** Shivali Kohli, Lilya Andrianova, Gabriella Margetts-Smith, Erica S Brady, Michael T Craig

**Affiliations:** Institute of Biomedical and Clinical Sciences, University of Exeter Medical School, Exeter EX4 4PS, UK; Institute of Neuroscience and Psychology, College of Medical, Veterinary and Life Sciences, University of Glasgow, Glasgow, G12 8QQ, UK

**Keywords:** Chemogenetics, thalamus, dementia

## Abstract

One of the main features of Alzheimer’s disease is the progressive loss of memory, likely due to pathological changes within brain regions such as the hippocampus and entorhinal cortex. These structures are embedded within the extended memory circuit, an interconnected set of brain regions that are essential for episodic memory. The anterior thalamic nuclei (ATN) and thalamic nucleus reuniens (NRe) are both extensively and reciprocally connected with these important memory regions, so we sought to test the hypothesis that chemogenetically-enhancing neurotransmission through NRe and ATN would ameliorate memory deficits in two mechanistically-distinct mouse models of Alzheimer’s disease. Using the hAPP-J20 mouse model of amyloidopathy and the Tg4510 mouse model of tauopathy, we carried out stereotaxic injections of viral vectors to transduce hM3Dq (Gq)_mCherry into NRe or the anterio-dorsal/anterio-ventral nuclei of ATN, using mCherry as a control. At nine months (hAPP-J20) or six months (Tg4510) of age, mice underwent a behaviour battery of open field (OF), novel object recognition (NOR) and radial arm maze (RAM), with DREADD agonist 21 administered 30min prior to each behaviour test. Tissue was collected post-behaviour to confirm injection site and virus expression. Both Tg4510 and hAPP-J20 mice show marked hyperactivity in the OF, significant deficits in recognition memory, and a significant impairment in spatial reference and spatial working memory. Unexpectedly, chemogenetic activation of ATN or NRe did not significantly improve spatial memory impairments or reduce the observed hyperactivity, although NRe activation did modestly rescue recognition memory in J20 mice. This may be due to compensation elsewhere within the memory circuit, or that the pathological changes are too far advanced for behaviour reversal.

## Introduction

Alzheimer’s disease is the most prevalent form of dementia, currently affecting an estimated 50 million people worldwide, with this number projected to triple by 2050^1^. The main features of Alzheimer’s disease include the progressive loss of memory and general cognitive decline, accompanied by extensive neurodegeneration in multiple brain regions, but particularly the hippocampus and entorhinal cortex^2^. Given their well-established roles in memory and spatial navigation, these two regions have received considerable focus in preclinical dementia research. However, these regions sit within an extended hippocampal memory system that comprises regions of the extended memory circuit, including prefrontal and retrosplenial cortices (RSC), and midline thalamic nuclei including nucleus reuniens (NRe) and the anterior thalamic nuclei (ATN)^3^. NRe forms reciprocal connections with medial prefrontal cortex (mPFC) and subiculum, and projects to hippocampal region CA1 and entorhinal cortex^4^. Similarly, the ATN are extensively interconnected with prefrontal, hippocampal and parahippocampal regions as well as the RSC and mammillary bodies^5^. Thus, the thalamus is well-placed to act as a communication hub between key areas of the brain’s extended memory circuit. Rodent studies show that NRe is required for goal-directed spatial navigation^6^ and that altering its activity is sufficient to impair working memory^7^. Similarly, lesions of ATN have long been known to disrupt hippocampal-dependent memory tasks^8^, with the ATN having roles in contextual fear memory^9^ and goal-directed behaviours^10^. Furthermore, recent advances suggest that both ATN and NRe could play a more general role in high-level information processing^11^. A recent review comparing function of the ATN with NRe concluded that the ATN support many aspects of hippocampal spatial encoding and performance while NRe aids a prefrontal, top-down control of cognition^12^. Altogether, the involvement of ATN and NRe in spatial memory is well-established, and a strong argument can be made for targeting these thalamic nuclei to treat memory disorders^3^.

The main pathological features for Alzheimer’s disease, beyond synaptic and neuronal loss, are the accumulation of amyloid-β plaques (Aβ) and neurofibrillary tangles of hyperphosphorylated microtubule-associated protein tau (referred to as ‘tau’ hereafter). Accumulations of neurofibrillary tangles have been observed in both the anterior-dorsal sub-region of the ATN and NRe in *post-mortem* human tissue^13^, and such pathological changes within the thalamus are associated with the memory loss seen in the early stages of Alzheimer’s disease progression in humans. This has not been widely studied in the myriad of animal models of Alzheimer’s disease available, although there are some reports of accumulations of both Aβ plaques and hyperphosphorylated tau in the thalamus of some animal models, reviewed by Aggleton and colleagues^14^. Nonetheless, irrespective of whether ATN and NRe show Alzheimer’s disease-related pathology in mouse models, they present a compelling target for therapeutic interventions in this disorder, given their role as a hub for communication throughout the extended memory circuit. Consequently, we sought to test this hypothesis using chemogenetic manipulation of ATN and NRe in preclinical models of dementia. We used two mechanistically-distinct mouse models: the hAPP-J20 amyloidopathy model, which over-expresses a form of hAPP that incorporates the Swedish and Indiana familial Alzheimer’s disease mutations^15^, and the Tg4510 tauopathy model that expresses the MAPT P301L mutation under the CaMKIIa promoter, limiting the expression of mutant tau to forebrain glutamatergic neurons^16,17^. Although degeneration in the Tg4510 mouse model is unlikely to be wholly attributed to neurofibrillary pathology^18^, it should not impede upon the ability to assess whether chemogenetic activation can improve spatial memory deficits in neurodegenerative disorders. In both mice strains, we expressed the hM3Dq excitatory Designer Receptor Exclusively Activated by Designer Drugs (DREADD)^19^ into ATN and / or NRe and assessed changes in learning and memory performance. Remarkably, despite their anatomical location as a ‘hub’ for transmission of spatial information in the brain, chemogenetic stimulation of these nuclei did not lead to improvements in spatial memory in either mouse model.

## Materials and Methods

### Animals

One-hundred-and-eighteen male and female hAPP-J20 mice (internally bred and maintained on a C57BL/6J genetic background) and seventy-eight male Tg4510 mice (kindly provided by Eli Lilly, UK) were housed in groups of two or three in individually ventilated cages with food and water available *ad libitum* until food restriction, where they were individually housed in the same cage type. Mice were maintained in a temperature (21 ± 2 °C) and humidity (45 ± 15%) controlled environment on a 12 h light-dark cycle (lights on at 06:20h), with all experiments performed in the light phase. All procedures were conducted in accordance with the Animals (Scientific Procedures) Act 1986 and received approval from the University of Exeter Local Ethical Committee Review Board.

In total, fourteen hAPP-J20 mice and seven Tg4510 mice were excluded from the study due to injection misplacement, whilst two hAPP-J20 and four Tg4510s were excluded from the RAM due to lack of habituation.

### Drugs

DREADD agonist 21 (Compound 21) dihydrochloride and clozapine-n-oxide (CNO) was purchased from Hello Bio (Avonmouth, UK), dissolved in 0.154 M saline, and subcutaneously administered at a volume of 1 mg/kg. The appropriate DREADD agonist and dose was selected based on current literature showing agonist 21 to have good bioavailability, pharmacokinetic properties, and brain penetrability^20^ without back-metabolism to clozapine, a common problem with other DREADD agonist, CNO^21^.

To establish that compound 21 at this dose shows no compounding effects on behaviour, six-month old Tg4510 mice (n=20) were tested on three occasions at weekly intervals in a novel object recognition (NOR) test following a subcutaneous (s.c.) injection of saline vehicle, compound 21 or CNO in a pseudo-random order to serve as their own control. All drugs were administered 30 min prior to each behaviour test; Supplementary Fig. 1.

### Stereotaxic surgery

Mice were anaesthetised with isoflurane in O_2_ and placed on a heat mat to maintain body temperature. Mice were placed in a stereotaxic frame and intracerebrally injected with 250 nl AAV8/2-hSyn-hM3D(Gq)_mCherry, 5.3×10^12^ vg/ml (vector v101-8, Viral Vector Facility, Neuroscience Centre Zurich, Switzerland) or AAV8/2-hSyn-mCherry 5.6×10^12^ vg/ml (vector v133-8, Viral Vector Facility, Neuroscience Centre Zurich, Switzerland) into the AD/AV thalamus (bilateral; AP −0.71, ML ± 0.8, DV 2.76 *from pia*) or NRe (unilateral; AP −0.8, ML 0.0, DV 3.8 *from pia*) respectively from Bregma (Paxinos and Watson) using a WPI syringe pump with a Hamilton syringe microinjector with 32 gauge, at a rate of 100 nl per minute. Representative images are shown in Supplementary Fig 2. Buprenorphine (0.03 mg/kg s.c.) was administered pre-surgery and Carprofen (1 mg/kg s.c.) was administered post-surgery. At the time of surgery hAPP-J20 mice were 7 – 7.5 months and Tg4510 mice were 4.5 months of age, respectively.

### Behaviour

Following the initial study determining the effects of compound 21 on the NOR behaviour task, all mice underwent a behaviour battery of open field, NOR and RAM. All tests were performed by an experienced observer. Tg4510 mice were tested at six months and hAPP-J20 mice were tested at nine months of age, respectively. The nine-month time point was chosen for hAPP-J20 mice, as they showed minimal behavioural deficits at six months (Supplementary Fig. 3). The experimenter was blind to mouse genotype and DREADD agonist administered in the pilot, and blind to mouse genotype, sex, and viral vector group in subsequent studies.

#### Open Field (OF)

To assess novel arena induced ambulatory activity mice were placed into individual Perspex boxes (40 x 40 x 45 cm) for 1 h and recorded using Ethovision video-tracking software (Noldus).

#### Novel Object Recognition (NOR)

To examine recognition memory mice underwent a two-trial object discrimination task. On the test day, mice were re-acclimatized to the OF box for 10 min, returned briefly to their home cage while two identical objects (red Lego tower, glass bottles or grey canisters) were placed in opposite corners. The mouse explored both objects for 10 min during the first familiarisation trial and the time (s) spent exploring each object was recorded from Ethovision video-tracking software (Noldus). Animals were returned to their home cage for an inter-trial interval (ITI) of 24 h before reintroduction to the cage with one original (familiar) object and one novel object. During the second trial, exploration of each object was once again recorded separately. The location of the novel object was varied in a pseudorandom order within groups. Exploratory behaviour was defined as sniffing, touching and direct attention to the object. Climbing on or chewing of the object was not counted. The Discrimination Index was calculated by the time spent exploring the novel object divided by the total time spent exploring both objects in the choice trial.

#### Radial arm maze (RAM)

To assess spatial working and reference memories, mice underwent the eight-arm RAM (Tracksys Ltd, UK). Each individual Perspex arm (38 x 7 x 15 cm, 45° from each other), originated from a central area used as the starting platform. 50% condensed milk was used as a food reward, and placed at the distal end of each arm. One-week prior mice were individually housed and food restricted to 90% bodyweight, which was maintained for the test duration. Two days before the test start, mice were habituated to the maze, allowed to explore freely, and consume the food reward. Successful habituation was deemed as consumption of all food rewards located in each arm, with the first reward consumed within 2 min of trial start. During the training sessions, four of the eight arms were baited for Tg4510 mice, and three of the eight baited for the hAPP-J20s. This was based on pilot data showing hAPP-J20 mice to have no significant impairment in the RAM using four-baited arms when tested at six months (Supplementary Fig. 6). The pattern of baited and un-baited arms remained unchanged throughout the entire procedure for each mouse. Each trial began by placing the mouse in the centre of the maze. Entry and re-entry into an un-baited arm were recorded as a reference memory (RM) error, and entry and re-entry into a previously visited arm was recorded as a working memory (WM) error. Trial ended when all food rewards had been consumed, to a maximum trial length of 5 min, and was conducted over 14 days.

### Tissue Collection

Post behaviour, mice were cervically dislocated with brains dissected and drop-fixed in 4% paraformaldehyde (Sigma-Aldrich, UK) to determine histological verification of correct stereotaxic injection placement and virus expression.

### Immunohistochemistry

Briefly, coronal sections of 30 μm thickness were obtained using a microtome (SM2010R, Leica, Germany). Free-floating sections were rinsed with phosphate-buffered saline (PBS) and endogenous peroxidase activity was blocked by incubation in 0.3% v/v hydrogen peroxide (Sigma, UK) in PBS for 15 min. Tissue sections were stained using the mouse-on-mouse (M.O.M.) immunodetection kit (VectorLabs; BMK-2202). Sections were blocked for 1 h in M.O.M.-blocking reagent followed by washing with PBS, and incubation with M.O.M.-diluent for 5 min, before incubation in mouse anti-MC1 (1:1000 dilution; a kind gift from Peter Davies) for 30 min at room temperature. Again, the sections were washed twice in PBS before incubation with M.O.M. Biotinylated IgG secondary antibody (1:250 dilution) for 10 min at room temperature. After reaction with avidin-biotin-horseradish peroxidase complex (Vectastain Elite ABC Standard Kit, Vector Laboratories) for 1 h, sections were developed using 0.04% 3, 3’-diaminobenzidine (DAB; HB0687, Hello-Bio Ltd, UK) and 0.05% nickel ammonium sulphate, in the presence of 0.04% v/v hydrogen peroxide in PBS. The reaction was stopped after sections were rinsed three times in PBS. Slices were mounted onto Superfrost Slides (#631-0108, VWR Int.) and dried overnight before dehydration using a graded series of alcohols and cleared using HistoClear (#Nat1334; Scientific Laboratory Supplies). Slides were cover-slipped using Histomount (#NAT1310; Scientific Laboratory Supplies) and left to dry for 2 days.

Bright-field images were taken as .TIFF files at a 10x magnification using a Nikon Eclipse 800 microscope using Qcapture (Qimaging) software.

### Statistical Analysis

Data were checked for normality and homogeneity of variance using Shapiro-Wilk’s and Levene’s tests respectively. Tg4510 mice behavioural data (all male) were analysed using either two way Analysis of Variance (ANOVA) tests – with main group factors viral vector and genotype – or three way mixed ANOVA with time, test day or object as a repeated measure factor. hAPP-J20 mice (male and female) behavioural data were analysed similarly; however tests were either three way ANOVA or four way mixed ANOVA, with sex as an additional main group factor for both. Statistical analysis was conducted using GraphPad Prism v9 (GraphPad Software Inc.) and IBM SPSS v25 software. Multiple comparison post-hoc tests were conducted where applicable. Data are presented as mean ± S.E.M. and significance α level set to *p* < 0.05.

### Data Availability

All raw data that supports the findings herein are available from the corresponding author upon reasonable request.

## Results

Following successful experiments to confirm compound 21 did not significantly alter baseline behaviour in the NOR paradigm in Tg4510 mice (Supplementary Fig. 1), mice underwent stereotaxic surgeries to inject viruses to deliver genes to express either hM3Dq_mCherry (with mCherry as a control), unilaterally or bilaterally, in the NRe or AD/AV thalamus, respectively. Representative images of excitatory DREADD expression in both NRe and AD/AV can be seen in Supplementary Fig. 2, and schematics showing the viral spread for all animals used in this study can be seen in Supplementary Fig. 3 (Tg4510 AD/AV experiment), Supplementary Fig. 4 (J20 AD/AV experiment) and Supplementary Fig. 5 (J20 NRe experiment).

### Chemogenetic excitation of the AD/AV thalamus does not ameliorate cognitive behaviour deficits in Tg4510 mice

During the OF, mice habituated to the arena with ambulatory counts progressively decreasing (*F_1, 27_* = *20.432*, *p* = *0.0001*; three way mixed ANOVA) over the 1 h time-course. There was a significant main effect of genotype in total distance travelled over the 1 h (*F_1,30_* = 19.470, *p* = *0.0001*), demonstrating that Tg4510 mice were significantly more active than WT controls, particularly at the 10 min (*p* < *0.01*), 40 min (*p* < *0.05*) and 50min (*p* < *0.05*) time points following Tukey post-hoc analysis. There were, however, no significant interactions reported between genotype or vector (p > 0.05).

Chemogenetic stimulation of AD/AV thalamus in Tg4510 mice failed to ameliorate this increased activity (*F_1,30_* = *2.579*, *p* = *0.120*), with Tg4510 mice expressing hM3Dq also showing markedly increased activity compared to wildtype controls, that was significant at the 40 min and 60 min time points (*p* < *0.05*; Tukey post-hoc); Fig. 1A and Fig. 1B. Indeed, chemogenetic activation of ATN in the Tg4510 may have exacerbated the hyperactive phenotype.

**Figure 1:**
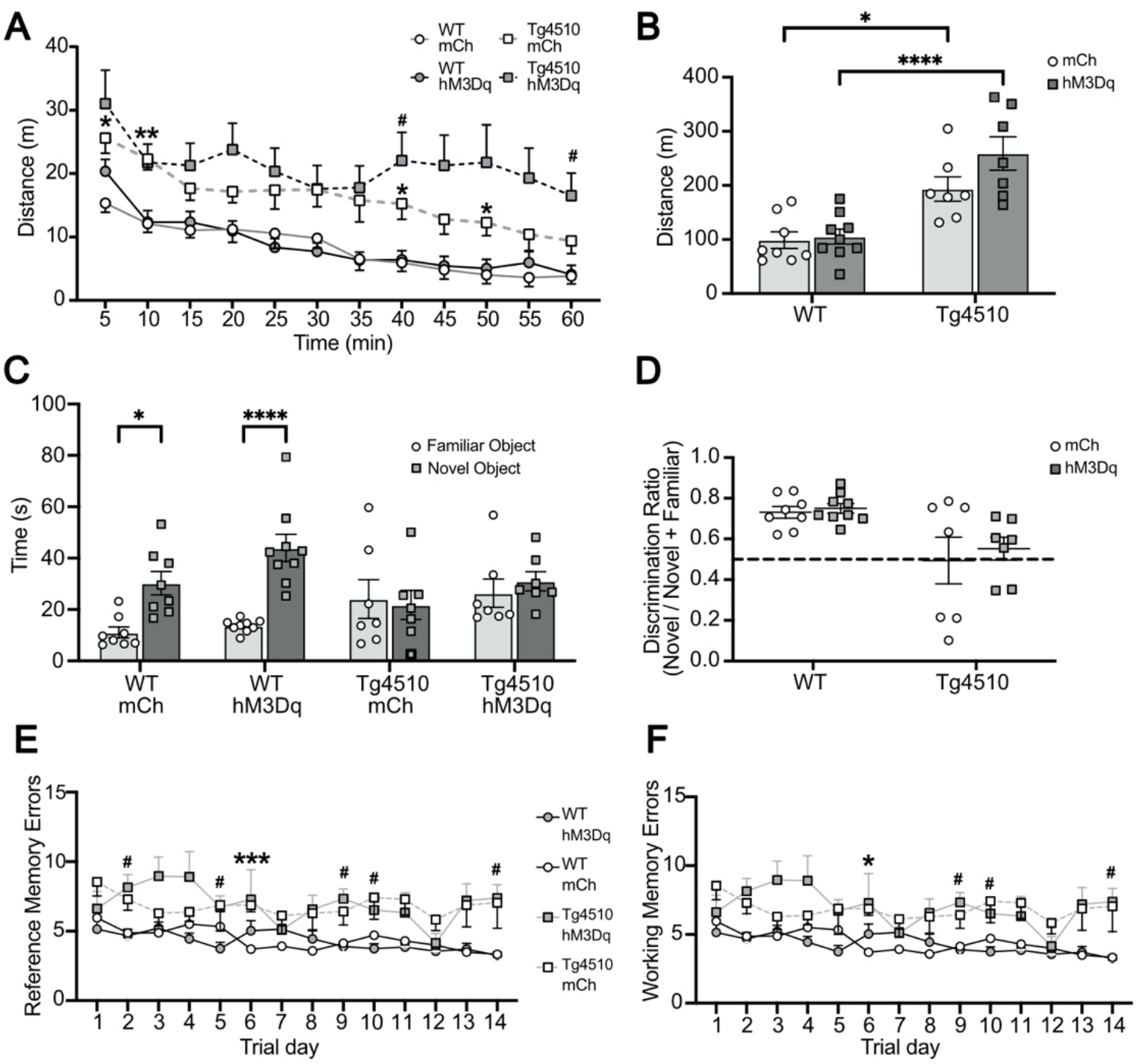
Effects of chemogenetic activation in the AD/AV thalamus in Tg4510 mice. Comparative differences (mean ± S.E.M.) in **A**, time-course of ambulatory activity (cumulative counts/5-min time-bins), \**p* < *0.05*, \*\**p* < *0.01* WT mCh vs Tg4510 mCh, #Tg4510 hM3Dq vs WT hM3Dq; Tukey post-hoc. **B**, total ambulatory count, **C**, time spent exploring familiar vs. novel object, Sidak post-hoc **D**, discrimination ratio in NOR after a 24h ITI, **E**, spatial reference and **F**, working memory errors in the RAM over 14 days in six month male Tg4510 mice and WT controls expressing excitatory DREADDs (hM3Dq; AAV8-hSyn-hM3Dq(Gq)_mCherry) or mCherry control (mCh; AAV8-hSyn-mCherry) in the AD/AV thalamus following subcutaneous administration of compound agonist 21 (1mg/kg) 30 min prior to each test. \**p* < *0.05*, \*\*\*\**p* < *0.0001*. AD/AV (anterio-dorsal/anterio-ventral thalamus), WT (wildtype), n=8 WT mCh, n=9 WT hM3Dq, n=7 Tg4510 mCh, n=7 Tg4510 hM3Dq.

In the NOR paradigm, a three-way mixed ANOVA revealed that WT mice spent significantly more time exploring the novel, as opposed to familiar, object following a 24h ITI (*F_1,27_* = *13.106*, *p* = *0.001*), which reached significance in both WT controls (*p* < *0.05*) and WT expressing excitatory DREADDs (*p* < *0.001*) following Sidak’s post-hoc test; Fig. 2C. However, male Tg4510 mice regardless of AAV vector, show deficits in recognition memory compared to WT controls as shown by the discrimination index (*F_1, 27_* = *10.860*, *p* = *0.03*), which reached post-hoc significance (*p* < *0.05*; Tukey post-hoc). This was not ameliorated by chemogenetic activation of the AD/AV thalamus via hM3Dq (*F_1, 27_* = *1.601*, *p* = *0.217*; Fig. 1D). Similarly, Tg4510 mice show robust deficits in working and learning memory relative to WT controls; chemogenetic enhancement of AD/AV activity failed to ameliorate neither reference (*F_1, 28_*=*65.247*, *p* = *0.0001*) nor working (*F_1, 28_*=*50.764*, *p* = *0.0001*) memory deficits; Fig. 1E and Fig. 1F. Due to limited availability of Tg4510 mice and the extensive neurodegeneration observed by the end of our behavioural battery (Supplementary Fig. 7), plus additional caveats associated with this mouse that became apparent midway through our study^18^, we did not test chemogenetic activation of NRe in these animals.

**Figure 2:**
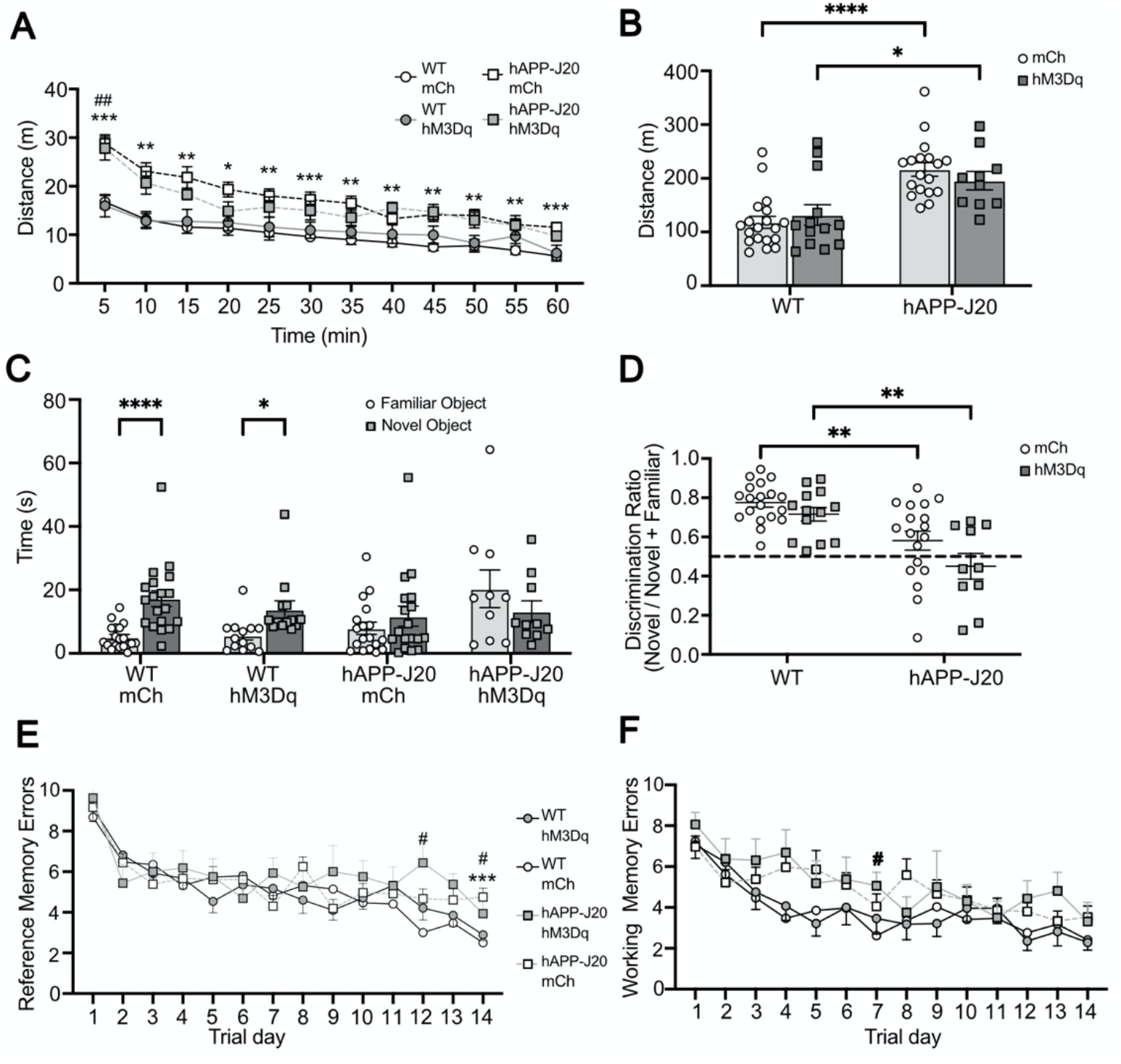
Effects of chemogenetic activation of the AD/AV thalamus in hAPP-J20 mice. Comparative differences (mean ± S.E.M.) in **A**, time-course of ambulatory activity (cumulative counts/5-min time-bins), \**p* < *0.05*, \*\**p* < *0.01*, \*\*\**p* < *0.001* WT mCh vs hAPP-J20 mCh, ##*p* < *0.01* WT mCh vs hAPP-J20 hM3Dq; Tukey post-hoc. **B**, total ambulatory count, Tukey post-hoc. **C**, time spent exploring familiar vs. novel object, Sidak post-hoc. **D**, discrimination ratio in NOR after a 24h ITI, Tukey post-hoc. **E**, spatial reference and **F**, working memory errors in the RAM over 14 days in nine month male and female hAPP-J20 mice and WT controls expressing excitatory DREADDs (hM3Dq; AAV8-hSyn-hM3Dq(Gq)_-mCherry) or mCherry control (mCh; AAV8-hSyn-mCherry) in the AD/AV thalamus following subcutaneous administration of compound 21 (1mg/kg) 30min prior to each test. No sex differences were found and so data is combined for representation. \**p* < *0.05*, \*\**p* < *0.01*, \*\*\**p* < *0.001*, \*\*\*\**p* < *0.0001*. AD/AV (anterio-dorsal/anterio-ventral thalamus), WT (wildtype), n=19 WT mCh, n=16 WT hM3Dq, n=19 hAPP-J20 mCh, n=15 hAPP-J20 hM3Dq.

### Chemogenetic excitation of the AD/AV thalamus does not ameliorate cognitive behaviour deficits in hAPP-J20 mice

During the OF, mice habituate to the arena, with ambulatory counts progressively decreasing over time (*F_1,52_* = *25.753*, *p* < *0.0001*; four-way mixed ANOVA) over the 1h time course. There was a significant main effect of genotype (*F_1,59_* = *25.823*, *p* < *0.0001*) in the total distance travelled, with hAPP-J20 mice exhibiting increased activity, with post-hoc significance for all time-points; Tukey post-hoc. There were however no significant main effects in total distance travelled between male and female mice (*F_1, 59_* = *0.27*, *p* = *0.869*) or following chemogenetic activation with excitatory DREADDs in the AD/AV thalamus (*F_1, 59_* = *0.0457*, *p* = *0.435*; Fig. 2A and Fig. 2B). There were also no significant interactions reported between genotype, sex or AAV administered (p > 0.05).

In the NOR paradigm, a four-way mixed ANOVA revealed that WT mice spent significantly more time exploring the novel object in the choice trial (*F_1,52_* = *8.065*, *p* < *0.05*) which also reached post-hoc significance for WT controls and those expressing hM3Dq in the AD/AV thalamus (*p* < *0.0001* and *p* < *0.05*, *respectively*; Sidak post-hoc). The ANOVA also revealed a significant interaction between object exploration and genotype (*F_1,52_* = *9.391*, *p* = *0.03*) and between object and viral vector (*F_1,52_* = *4.024*, *p* = *0.05*) but no significant interaction between object and sex (p > 0.05).No sex differences were observed in either WT or hAPP-J20 mice (*F_1, 59_* = *1.390*, *p* = *0.244*), but hAPP-J20 mice show impaired recognition memory as shown by the discrimination index (*F_1, 59_* = 22.792, *p* < *0.0001*), which was not rescued by chemogenetic activation within the AD/AV thalamus (*F_1, 59_* = *2.862*, *p* = *0.097*; Fig. 2C and 2D).

Within the RAM, hAPP-J20 mice also show impaired spatial reference (*F_1, 49_* = *4.242*, *p* = *0.045*) and spatial working (*F_1, 49_* = *10.766*, *p* = *0.002*) memory deficits compared to WT controls following a four-way mixed ANOVA. Similarly, no sex differences were seen in learning for either reference (*F_1, 49_* = *1.288*, *p* = *0.262*) or working memory (*F_1, 49_* = *1.155*, *p* = *0.288*) performance, nor a significant main interaction between genotype, sex or AAV vector (p > 0.05.). Chemogenetic activation of the AD/AV thalamus did not influence learning in either WT mice, nor improve impaired learning deficits in hAPP-J20 mice for either reference (*F_1,49_* = *0.364*, *p* = *0.549*), or spatial working (*F_1,49_* = *0.004*, *p* = *0.953*) memory performance; Fig. 2E and 2F).

### Chemogenetic excitation of NRe does not ameliorate cognitive behaviour deficits in hAPP-J20 mice

During the OF, mice habituate to the arena, with distance travelled progressively decreasing over time (*F_1,51_* = *73.576*, *p* < *0.0001*; four-way mixed ANOVA) over the 1h time course. There was a significant main effect of genotype (*F_1,58_* = *73.577*, *p* < *0.0001*) in the total distance travelled, with hAPP-J20 mice exhibiting increased activity, with post-hoc significance reached for all time points; Tukey post-hoc; Fig. 3A). There were however no significant main effects in total distance travelled between male and female mice (*F_1, 58_* = *1.828*, *p* = *0.182*) nor following chemogenetic activation with excitatory DREADDs in NRe (*F_1, 58_* = *0.619*, *p* = *0.435*; Fig. 3B). There were also no significant interactions reported between genotype, sex or AAV administered (p > 0.05).

**Figure 3:**
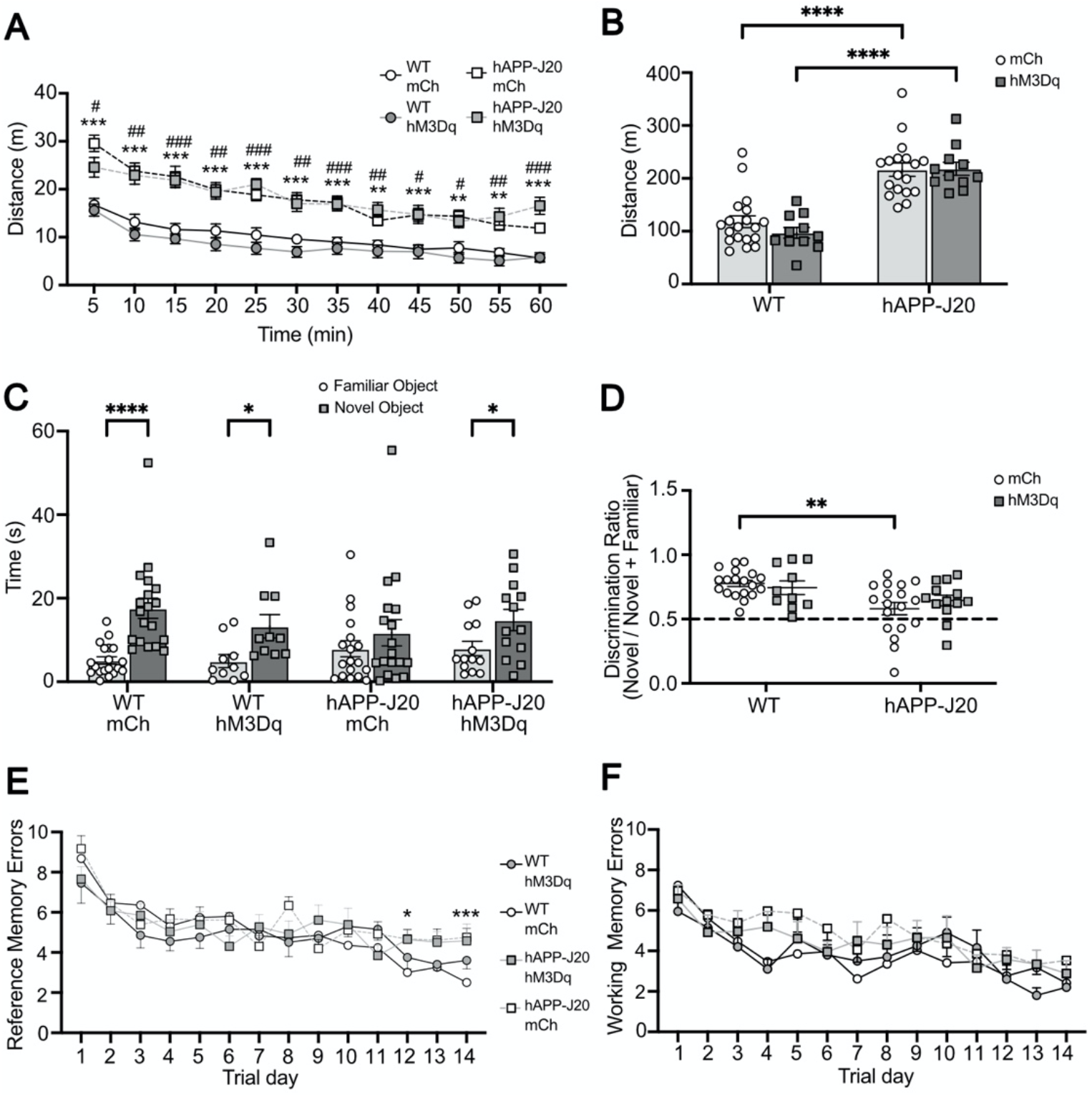
Effects of chemogenetic activation of NRe in hAPP-J20 mice. Comparative differences (mean ± S.E.M.) in **A**, time-course of ambulatory activity (cumulative counts/5-min time-bins), \*\**p* < *0.01*, \*\*\**p* < *0.001* WT mCh vs hAPP-J20 mCh, #*p* < *0.05*, ##*p* < *0.01*, ###*p* < *0.001* WT mCh vs hAPP-J20 hM3Dq; Tukey post-hoc. **B**, total ambulatory count, Tukey post-hoc. **C**, time spent exploring familiar vs. novel object, Sidak post-hoc. **D**, discrimination ratio in NOR after a 24h ITI, Tukey post-hoc. **E**, spatial reference and **F**, working memory errors in the RAM over 14 days in nine month male and female hAPP-J20 mice and WT controls expressing excitatory DREADDs (hM3Dq; AAV8-hSyn-hM3Dq(Gq)_-mCherry) or mCherry control (mCh; AAV8-hSyn-mCherry) in the NRe following subcutaneous administration of compound 21 (1mg/kg) 30min prior to each test. No sex differences were found and so data is combined for representation. \**p* < *0.05*, \*\**p* < *0.01*, \*\*\**p* < *0.001*, \*\*\*\**p* < *0.0001*. NRe (Nucleus Reuniens), WT (Wildtype), n=19 WT mCh, n=15 WT hM3Dq, n=19 hAPP-J20 mCh, n=15 hAPP-J20 hM3Dq.

In the NOR paradigm, a four-way mixed ANOVA revealed that WT mice spent significantly more time exploring the novel object in the choice trial (*F_1,51_* = *36.368*, *p* = *0.001*) which also reached post-hoc significance (*p* < *0.0001* and *p* < *0.05*; Sidak post-hoc) for WT controls and those expressing excitatory DREADDs in the AD/AV thalamus; Fig. 3C. The ANOVA also revealed a significant interaction between object exploration and genotype (*F_1,52_* = *5.731*, *p* = *0.02*) and between object and viral vector (*F_1,52_* = *3.349*, *p* = *0.079*) but no significant interaction between object and sex (p > 0.05). No sex differences were observed in either WT or hAPP-J20 mice (*F_1, 58_* = *2.806*, *p* = *0.100*), but hAPP-J20 mice show impaired recognition memory as shown by the discrimination index (*F_1, 58_* = *14.269*, *p* < *0.0001*). In the choice trial, compound 21-treated hM3Dq-injected hAPP-J20 mice show increase exploration for the novel as opposed to familiar object, reaching post-hoc significance (*p* < *0.05*; Sidak post-hoc) but this effect was insufficiently large to see a significant main effect of chemogenetic activation in NRe (*F_1,58_* = *0.913*, *p* = *0.344*; Fig. 3D). Nonetheless, this modest rescue of recognition memory with chemogenetic stimulation of NRe in J20 mice is consistent with a recent report of NRe playing a role in recognition memory^22,23^.

Within the RAM, hAPP-J20 mice also show impaired spatial reference (*F_1, 50_* = *4.535*, *p* = *0.038*) and spatial working (*F_1, 50_*=*8.002*, *p* = *0.007*) memory deficits compared to WT controls following a four way mixed ANOVA. Similarly, no sex differences were seen in learning for either reference (*F_1, 50_* = *1.615*, *p* = *0.210*) or working memory (*F_1, 50_* = *2.854*, *p* = *1.288*) performance, nor a significant main interaction between genotype, sex or AAV vector (*p* > 0.05). Chemogenetic activation of thalamic nucleus reuniens did not influence learning in either WT mice, nor improve memory deficits in hAPP-J20 mice for either reference (*F_1,50_* = *0.458*, *p* = *0.502*),or spatial working (*F_1,50_* = *0.782*, *p* = *0.381*) memory performance; Fig. 3E and 3F.

## Discussion

This study set out to test the hypothesis that spatial memory impairments in mouse models of amyloidopathy and tauopathy could be improved by providing exogenous chemogenetic activation using DREADDs to the two midline thalamic nuclei known to act as a conduit for communication throughout the hippocampal-entorhinal-prefrontal network^12^. We found that chemogenetic activation of neurons in these nuclei neither ameliorated spatial memory deficits in transgenic mice nor disrupted memory in WT animals. While this result was somewhat unexpected, it provides evidence that these thalamic systems are remarkably resistant to perturbation in both wildtype mice and those that model dementia.

### The role of ATN in spatial memory

Much of our understanding of the role of ATN in spatial memory come from lesion studies, as has been reviewed by Aggleton and Nelson^8^; these rodent studies show that the ATN play an integral role in processing information from local and distant circuits during memory formation by coordinating and synchronising activity across hippocampal and cortical regions. This function has also recently been demonstrated in human epilepsy patients undergoing combined intrathalamic and scalp EEG recordings^24^. Lesions in the ATN have been shown to reduce activity within the RSC; an area also implicated in spatial and contextual memory formation^25,26^. In addition to lesion studies, it was recently reported that chemogenetic inhibition of ATN is sufficient to impair spatial memory in rats^27^; this study found that this effect could also be replicated by inhibiting the dorsal subiculum projections to ATN.

One possible explanation for the failure to find evidence of an effect in our experiment could be that pathology in hippocampal regions upstream of ATN could be too advanced for manipulating ATN to then have a positive effect in rescuing spatial memory. This could certainly be true of the Tg4510 mouse line, which displays substantial neurodegeneration throughout the entire extended memory circuit by six to seven months (Supplementary Fig. 5). While we are not aware of any published study that has used chemogenetic or optogenetic stimulation of ATN to rescue memory deficits, the connectivity of ATN and lesion studies suggest this should be a successful strategy, as also discussed by Barnett and colleagues^28^. Electrical stimulation of ATN in rats, when combined with corticosterone delivery, significantly improves performance in the delayed nonmatch to sample T maze task with a delay of 60 s, but significantly impairs performance at longer latencies^29^. In humans, long-term deep brain stimulation of ATN in epilepsy patients was found to improve performance on memory tasks^30^. We employed a chemogenetic strategy as we predicted that optogenetic stimulation in a massively interconnected network would disrupt memory, whereas chemogenetic stimulation would serve to amplify endogenous signalling. However, the success of electrical stimulation suggests that perhaps a stronger drive of ATN neurons is required to improve memory, so optogenetic stimulation may prove more fruitful at rescuing memory deficits. Indeed, a recent study by Barnett and colleagues found that optogenetic stimulation of ATN could reverse memory deficits in rats with lesions of the mammillothalamic tract^31^. Of relevance to our study was their observation that optogenetic stimulation, applied at theta-burst frequency, rescued memory deficits when the timing of the burst was exogenously applied, but not when the timing was coupling to the rat’s endogenous hippocampal theta rhythm. This suggests that memory impairments can indeed be rescued by targeting the ATN, but the timing of stimulation is crucial; it may be that the chemogenetic approach employed in our study lacks the temporal precision required to rescue memory deficits.

### The role of NRe in spatial and object memory

A similar argument may be made for NRe: acting as a hub for frontal and hippocampal interactions, NRe could also be an attractive target for treating memory impairments. Indeed, NRe contains head-direction cells^32^ and lesions in this area also contribute to impaired spatial learning performance in the Morris Water Maze^33,34^. Transient inactivation of the NRe via GABAA agonist muscimol in rats has been shown to produce deficits in a T-maze task^35^ via regulation of hippocampal-PFC synchrony^36^, and optogenetic inhibition of NRe afferents in CA1 is sufficient to impair goal-directed behaviour in mice^6^. In contrast to ATN, where inactivation disrupts spatial memory^27^, optogenetic stimulation in NRe has been reported to disrupt spatial memory^7^. Interestingly, while ATN tends to interact more with the dorsal hippocampus, NRe preferentially (but not exclusively) engages with ventral hippocampal regions, suggesting NRe may associate more with object-based, rather than spatial, information^12^. Our findings are consistent with this perspective: chemogenetic activation of NRe failed to alter performance in the eight-arm radial maze in either WT or hAPP-J20 mice in our study. The hAPP-J20 mice in our study also displayed impaired novel object recognition; while our overall analysis failed to find strong evidence of an effect of chemogenetic manipulation on novel object recognition, *post hoc* multiple comparisons (Fig. 4C) provided evidence of a modest improvement in novel object recognition in the compound 21-treated over vehicle-treated hAPP-J20 mice. Recent work has shown that NRe inactivation impairs aspects of recognition memory^22,23^, so the improvement observed in our study is consistent with these findings, and serves to act as a crucial control demonstrating that we were achieving sufficient activation using chemogenetic tools to effect changes in behaviour.

### Midline thalamus as a therapeutic target in dementia?

We used two mechanistically-distinct mouse models of Alzheimer’s disease in an attempt to determine whether the midline thalamus could serve as a novel therapeutic target to treat memory impairment. Our results indicate that acute treatment with excitatory DREADD agonists was unable to attenuate behaviour deficits in two different models despite robustly activating thalamic neurons. However, this does not mean that future progress in this area is outwith the realm possibility. It may be that more specific targeting of individual projections through the thalamus is needed, as opposed to activating all outputs from the thalamic nuclei to their target regions (many of which are also interconnected through direct projections). Recent work by Nelson and colleagues^27^ suggests that selectively activating the dorsal subiculum to ATN projection could yield positive results. This could be achieved using an intersectional strategy, by delivering Cre-recombinase into a retrograde virus into ATN and using a Cre-dependent effector in dorsal subiculum.

Although both mouse models used in our study have been well characterised, further work is required to understand which circuit mechanisms are impaired, and thus amenable to being targeted by chemogenetic tools. For example, extracellular field recordings in the CA1/subiculum of hAPP-J20 mice show reduced theta-gamma cross-frequency coupling, along with reduced intrinsic excitability of parvalbumin interneurons^37^. As inhibitory parvalbumin cells play an important role in coordinating hippocampal theta and gamma oscillations^38^, and the coupling of these oscillations play a pivotal role in spatial learning^39^, using chemogenetic approaches to instead recruit GABAergic populations within local hippocampal circuitry, could improve memory. Indeed, previously we found that perturbation of excitatory inputs onto parvalbumin interneurons exacerbated deficits in gamma oscillations in the APPswe/PS1DE mouse model^40^, and our recent study using in the APP^NL-G-F^ knock-in mouse line^41^ found that subtle changes in hippocampal network oscillations during natural sleep were associated with an increase in synchrony of hippocampal PV interneurons^42^.

Another potential explanation for our failure to observe a positive result could be that any manipulation to thalamic excitability needs to occur before the onset of behavioural changes in the mouse models that we used. In particular, the Tg4510 mouse model displayed severe widespread neurodegeneration at the time point used (Supplementary Fig. 5). Long-term ATN stimulation improved memory in human epilepsy patients^30^ so it may be that chronic activation of ATN, perhaps via inclusion of DREADD agonists in the rodent chow, is needed to reverse dementia-related dysfunction in the extended memory circuit. The thalamus shows pathological changes within Alzheimer’s disease, and mouse models of dementia suggest similar pathological changes^14^. This may therefore be a limitation of currently available mouse models, and modelling robust thalamic pathologies may require combined interactions between amyloid and tau pathologies, and even glial interactions that are not currently captured. For example, post-mortem tissue from Down syndrome patients diagnosed with dementia, revealed fewer glial cells, in addition to neuronal loss and reduced volume in the ATN (incorporating the anterior-ventral and anterior-medial nuclei of the thalamus), compared to age-matched controls^43^. Glial loss in the ATN of hAPP-J20 mice has not been well characterised, however increased GFAP-positive astrocytes are observed in the hippocampus, alongside neuronal loss in 9-month-old mice^44^. Furthermore, there is scarce literature investigating thalamic projections to and from key regions within the extended memory circuit in hAPP-J20 mice, and so synaptic functions in the ATN and its connections may not be sufficiently disrupted before the appearance of cognitive impairment. If this is true, it may be of benefit to instead reduce thalamic excitability in the same brain regions by the use of inhibitory DREADDs, and determine whether this would influence cognitive behaviours in either WT or transgenic mice. It may also be of interest to use modern approaches combining wireless *in vivo* recording technology alongside behaviour and decisionmaking tasks. This would provide more insight as to how circuit manipulation via specific sub-regions can influence behavioural outcomes and neuronal firing in real-time.

### Potential off-target effects of compound 21?

A recent study reported that the dose of compound 21 used in our study (1 mg/kg) can cause sex-dependent off-target effects in the firing of neurons in subtantia nigra^45^. It should be noted that this study used intraperitoneal injection in rats, while our study used subcutaneous injection in mice like our present study. However, the report by Goutaudier and colleagues^45^ is consistent with earlier radioligand binding data showing that compound 21 has affinity for both D1 and D2 dopamine receptors^46^. While the dose of compound 21 that we employed in our study was at the upper end of those used in these two studies, it was found to have little or no nonspecific effects on behaviour^46^. Additionally, we carried out a pilot study to test the effects of CNO and compound 21 on the locomotor activity and novel object recognition memory in surgical naïve mice and found no effect of either agonist on behaviour (Supplementary Fig. 1). This, in addition with our mCherry vs mCherry-hM3Dq controls, makes us confident that any off-target effects of compound 21 had no effect on the outcome of our study.

### Conclusions and Future Directions

Surprisingly, we found that chemogenetic activation of ATN was unable to reverse spatial memory deficits in mouse models of dementia, nor was it sufficient to disrupt spatial memory in wildtype mice. The thalamus receives key inputs from the hippocampal formation and mammillary bodies, with cortical and hippocampal projections modulating neuronal activity in the anterior dorsal ATN region^47^. Excitatory DREADDs may be activating cells within the ATN and NRe, but this does not necessarily spread to wider cortical and subcortical circuits involved in spatial navigation and memory. It may also be that widespread activation of a strongly interconnected circuit was too broad to have the desired effect. Although projections to and from the ATN and NRe play a role in spatial working memory, improving neurotransmission through these central hubs was not sufficient in attenuating a strong behavioural impairment. Potentially, this style of stimulation or increased excitability needs to be introduced at a younger age in our mouse models, before strong pathological and behavioural changes are observed. Future studies will need to record neuronal network activity during such behaviour tasks, to provide more insight into how excitatory DREADDs may influence circuit-level neuronal oscillatory activity, and how these changes can influence behavioural outcomes in mice.

## Abbreviations

AD/AV: (Anterior-dorsal/Anterior-ventral)
DREADDs: (Designer Receptors Exclusively Activated by Designer Drugs)
NRe: (Nucleus Reuniens)
WT: (Wildtype)

## Acknowledgments

MTC and SK conceived and designed the study, and drafted the manuscript. SK carried out behavioural and immunohistochemistry experiments. SK, LA, GMS, EB and MTC optimised and carried out stereotaxic surgeries and viral injections. SK, LA, GMS and EB maintained animal colonies. All authors discussed the results and critically revised the manuscript for intellectual content. Tg4510 mice were kindly provided by Eli Lilly (UK), and the MC1 antibody was kindly provided by Peter Davies, Albert Einstein College of Medicine (USA). Finally, we are grateful to the anonymous peer reviewers who provided constructive critical feedback that helped us to significantly strengthen our manuscript.

