## Supplementary figures for "Chemogenetic activation of midline thalamic nuclei fails to ameliorate memory deficits in two mouse models of Alzheimer’s disease"

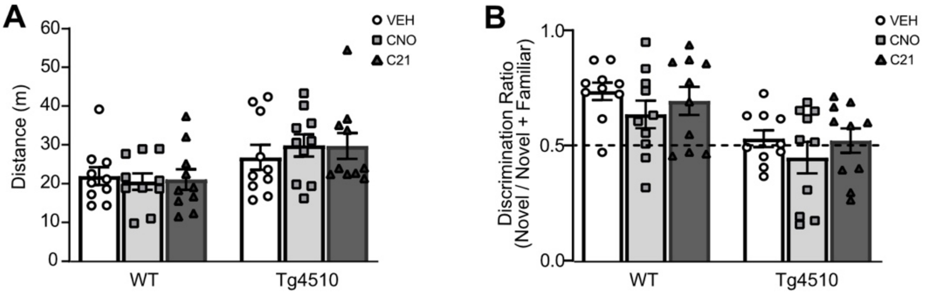


**Supplementary figure 1: Effects of DREADD agonists on drug- and surgery-naïve male Tg4510 mice.**
**A,** Tg4510 mice have significantly higher ambulatory activity in the open field, with a greater distance travelled in 10min ($F_{1,54} = 11.21$, $p = 0.0015$), that is not influenced by DREADD agonists ($p > 0.05$; two-way ANOVA). **B,** Tg4510 mice have significantly impaired recognition memory following a 24h ITI compared to WT controls ($F_{1,54} = 18.20$, $p = 0.0001$) that is not influenced by DREADD agonists ($p > 0.05$.; two-way ANOVA). Graphs show comparative differences (mean ± S.E.M.). Behaviour conducted after s.c. administration of saline (VEH), or DREADD agonists Clozapine-N-Oxide (CNO; 1mg/kg) or Compound 21 (C21; 1mg/kg) 30min prior. n=10 WT (Wildtype), n=10 Tg4510.

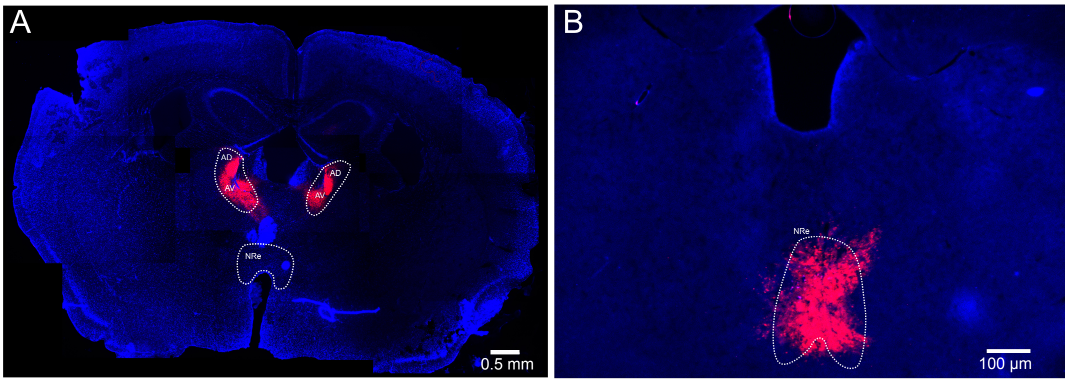


**Supplementary figure 2: Representative images of stereotaxic injections A;** Anterio-dorsal/Anterio-ventral (AD/AV) thalamus, and **B;** Nucleus Reuniens (NRe).

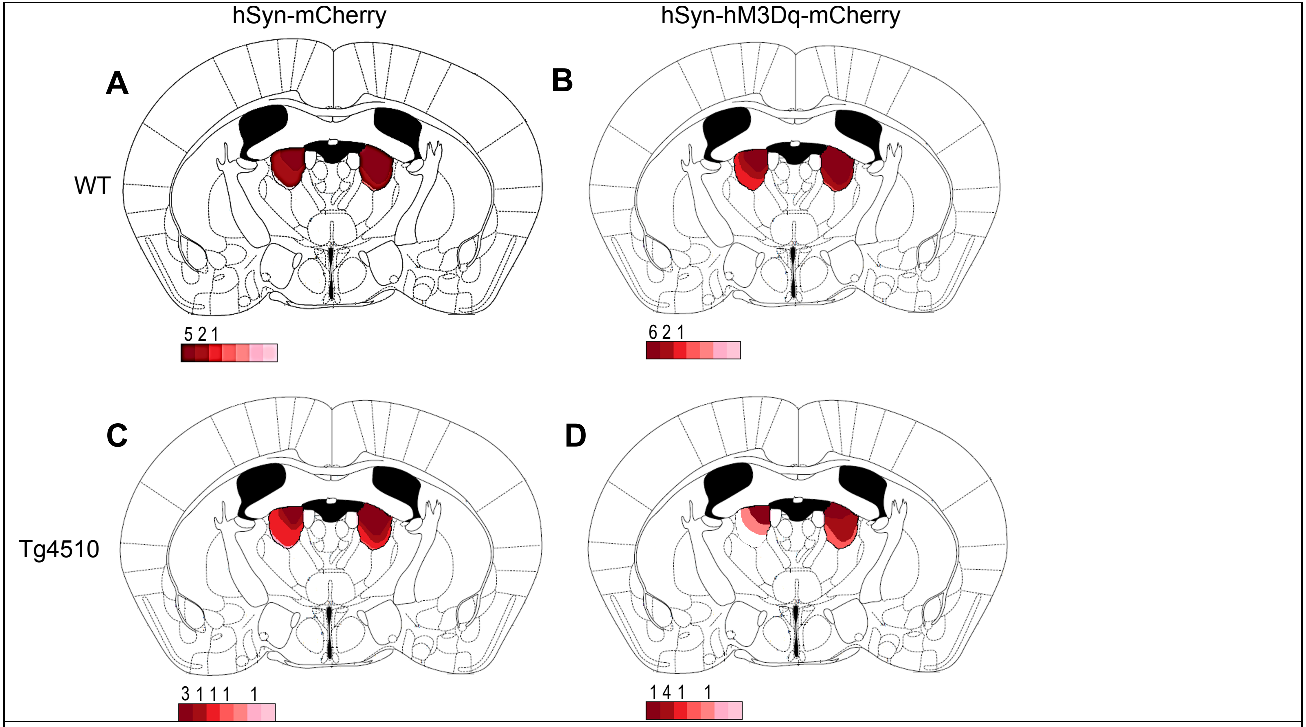


**Supplementary figure 3: outline of viral spread in all Tg4510 mice and WT controls in AD/AV experiment.** An approximated schematic of viral spread for all J20 mice used within this study is also shown for **A,** wildtype mice injected with -mCherry (n=8); **B,** wildtype mice injected with hM3Dq-mCherry (n=9); **C**, Tg4510 mice injected with -mCherry (n=7); and **D**, Tg4510 mice injected with hM3Dq-mCherry (n=7).

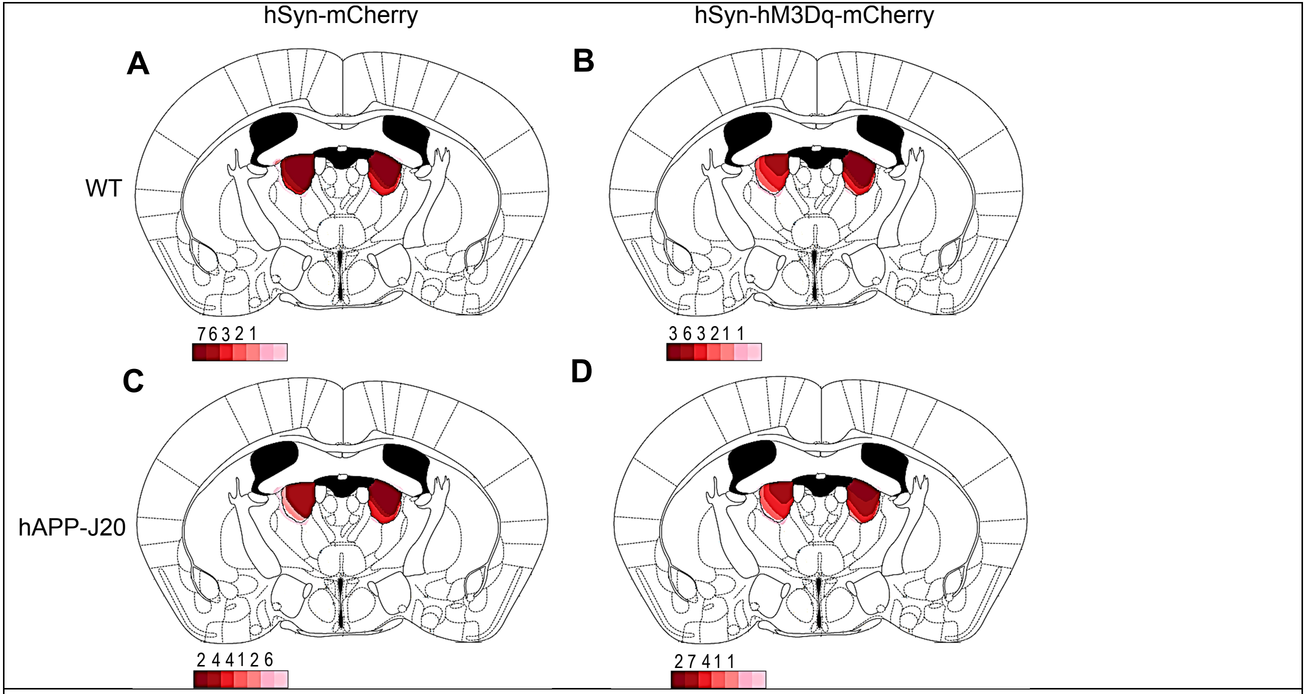


**Supplementary figure 4: outline of viral spread in all hAPP-J20 mice and WT controls in AD/AV experiment.** An approximated schematic of viral spread for all J20 mice used within this study is also shown for **A**, wildtype mice injected with -mCherry (n=19); **B**, wildtype mice injected with hM3Dq-mCherry (n=16); **C**, hAPP-J20 mice injected with -mCherry (n=19); and **D**, hAPP-J20 mice injected with hM3Dq-mCherry (n=15).

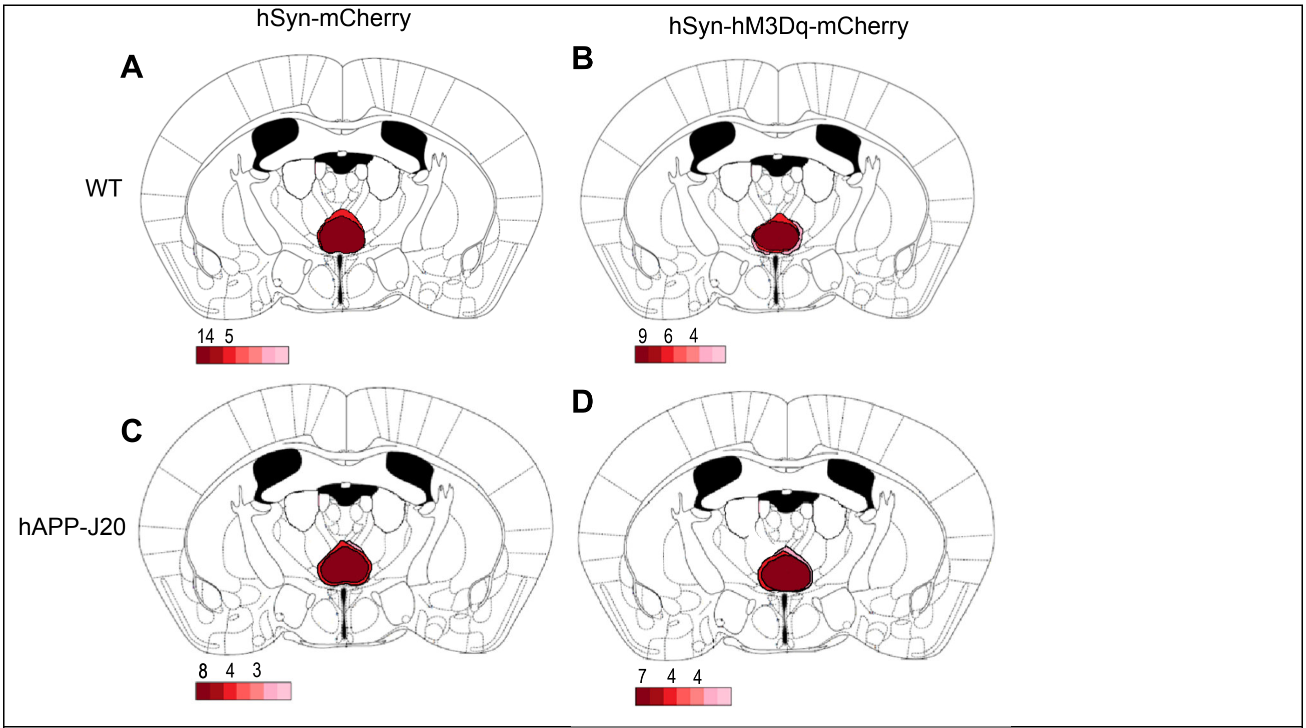


**Supplementary figure 5: outline of viral spread in all hAPP-J20 mice and WT controls in NRe experiment.** An approximated schematic of viral spread for all J20 mice used within this study is also shown for **A**, wildtype mice injected with -mCherry (n=19); **B**, wildtype mice injected with hM3Dq-mCherry (n=15); **C**, hAPP-J20 mice injected with -mCherry (n=19); and D, hAPP-J20 mice injected with hM3Dq-mCherry (n=15).

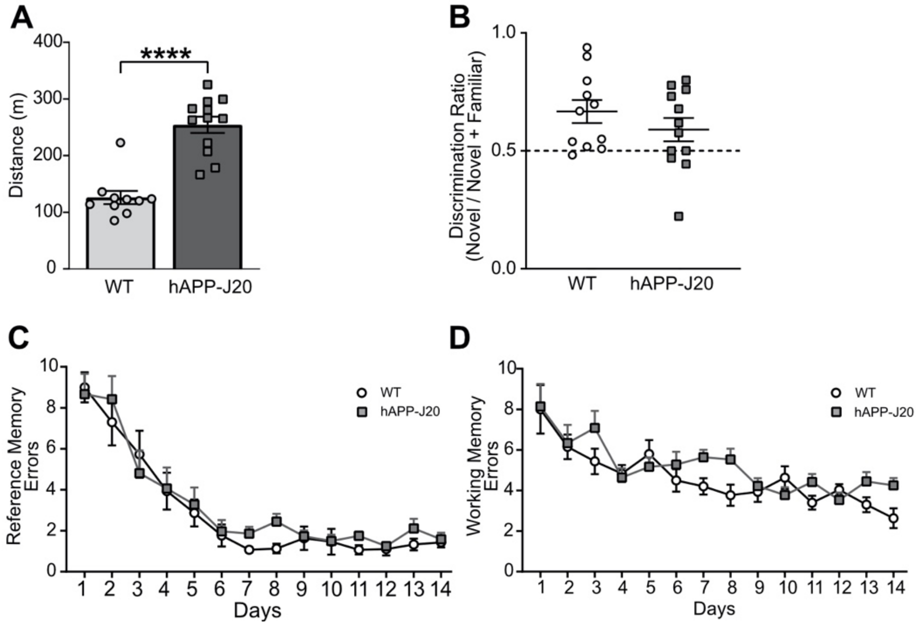


**Supplementary figure 6: Cognitive behaviour changes in six-month old hAPP-J20 mice.**
**A,** hAPP-J20 mice show significantly more activity (mean ± S.E.M. = 234.0 ± 13.18m) in total distance travelled in 1h compared to WT controls (mean ± S.E.M. = 116.3 ± 10.73m; $t_{20} = 6.738$, $p < 0.0001$ unpaired Student's *t*-test). **B,** hAPP-J20 mice show no significant impairment in the NOR ($t_{20} = 1.347$, $p > 0.05$. unpaired Student's *t*-test) after a 24h ITI. In the RAM, a two-way RM ANOVA of C, spatial reference ($F_{1,20} = 1.6389$, $p > 0.05$) and D, working memory ($F_{1,20} = 3.270$, $p = 0.0856$) errors revealed no significant effects of genotype. Graphs show comparative differences (mean ± S.E.M.). WT (Wildtype), n=10 WT, n=12 hAPP-J20.

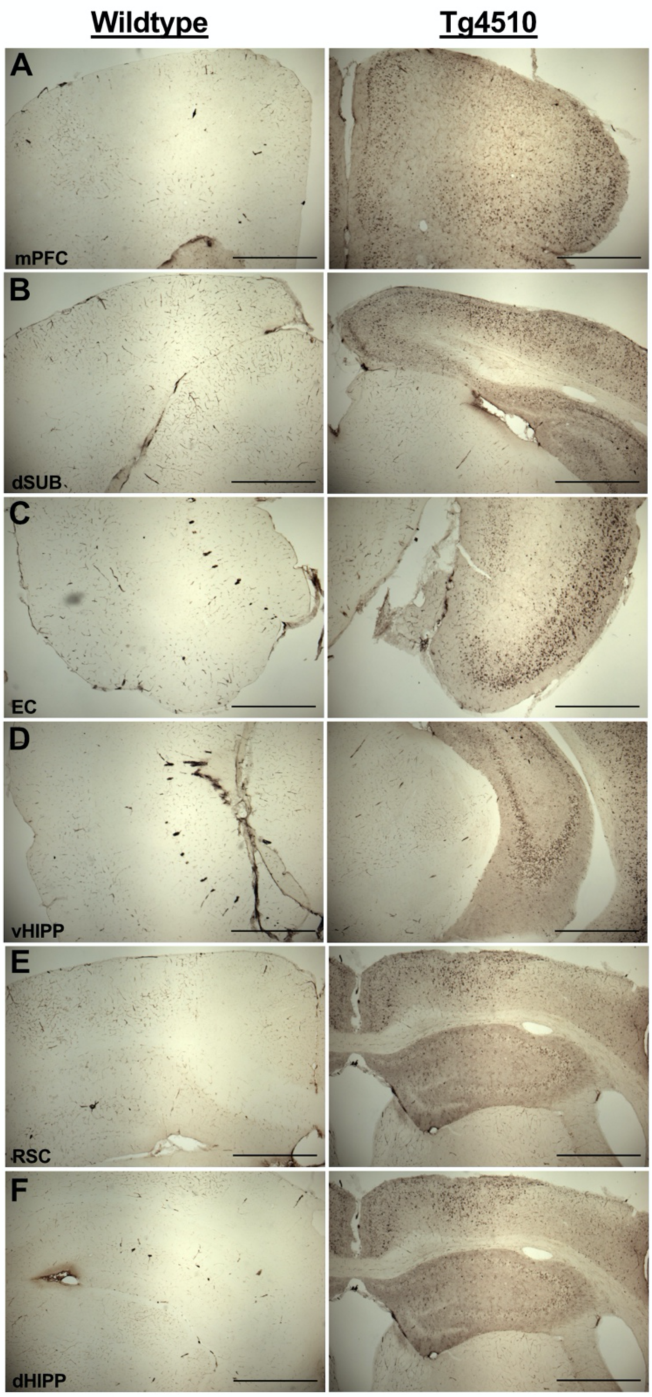


**Supplementary figure 7: Tg4510 mice show substantial degeneration at seven months.** Representative x10 magnification images of seven-month old Wildtype and Tg4510 mice showing hyperphosphorylated tau (MC1 antibody) in the **A;** medial prefrontal cortex (mPFC), **B;** dorsal subiculum (dSUB), **C;** entorhinal cortex (EC), **D;** ventral hippocampus (vHIPP), **E;** retrosplenial cortex (RSC) and **F;** dorsal hippocampus (dHIPP) in tissue stained for monocolonal antibody MC-1. Scale bar = 100µm.
